# Distinct microbial and immune niches of the human colon

**DOI:** 10.1101/2019.12.12.871657

**Authors:** KR James, T Gomes, R Elmentaite, N Kumar, EL Gulliver, HW King, MD Stares, BR Bareham, JR Ferdinand, VN Petrova, K Polanski, SC Forster, LB Jarvis, O Suchanek, S Howlett, LK James, JL Jones, KB Meyer, MR Clatworthy, K Saeb-Parsy, TD Lawley, SA Teichmann

## Abstract

Gastrointestinal microbiota and immune cells interact closely and display regional specificity, but little is known about how these communities differ with location. Here, we simultaneously assess microbiota and single immune cells across the healthy, adult human colon, with paired characterisation of immune cells in the mesenteric lymph nodes, to delineate colonic immune niches at steady-state. We describe distinct T helper cell activation and migration profiles along the colon and characterise the transcriptional adaptation trajectory of T regulatory cells between lymphoid tissue and colon. Finally, we show increasing B cell accumulation, clonal expansion and mutational frequency from caecum to sigmoid colon, and link this to the increasing number of reactive bacterial species.

## Introduction

The colon, as a barrier tissue, represents a unique immune environment where immune cells display tolerance towards a diverse community of microbes - collectively known as the microbiome. The microbiome is critical for many aspects of health and an imbalance of commensals and pathogenic microbes is linked with many disease states ^1^. Thus, understanding what constitutes a healthy, homeostatic relationship between host immune cells and the microbiome of the human colon is of critical importance. At a coarse-grained level, intestinal microbiota and immunity have been studied extensively. Here, we provide an in-depth map of proximal-to-distal changes in the colonic microbiome and host immune cells.

The composition of the microbiota at any location in the intestines is determined by the availability of nutrients, oxygen and the transit rate of luminal content, and as such, is spatially distinct ^2^. Regional differences are most evident when comparing the small intestine and distal colon in humans or other mammals ^3,4^. Within the colon, early studies identified changes from proximal (including the caecum, ascending and transverse colon) to distal colon (comprising the descending and sigmoid colon connecting to the rectum). Changes were evident with respect to the growth rates and activities of some bacteria species, which were dependent on nutrient availability ^4^. A proximal-to-distal gradient of bacterial diversity in which the number of operational taxonomic units (OTU) increases has been reported ^5^.

The intestinal immune system has a symbiotic relationship with the microbiome and is central to the maintenance of epithelial barrier integrity. The lamina propria and associated lymphoid tissues contain one of the largest and most diverse communities of immune cells - including both lymphocytes and myeloid cells ^6^. There is marked regional variation in immune cells along the gastrointestinal tract, with T helper (Th) 17 cells decreasing in number from duodenum to colon, and T regulatory (Treg) cells being highest in the colon ^7^. Whether there is further complexity, with distinct immunological niches, within the colon remains to be elucidated. Immune cells can respond to environmental cues including the microbiota. Mouse studies have demonstrated that specific bacterial species can fine-tune intestinal immune responses, including Th17 ^8,9^, Treg ^10^, or Th1 ^11^ polarisation and function ^8,12^ and B cell activation ^13^. However, the extent to which there is regional variation in the mucosal microbiome within an individual, and how this might influence local immune cell niches along the colon, has not been investigated to date.

Here, we catalogued the mucosal microbiome in different regions of the human colon, a gastrointestinal organ with the most diverse and dense microbiome content and region-restricted disease states ^14^. In parallel, we applied single-cell RNA-seq (scRNA-seq) to make a census of steady-state immune cell populations in the adjacent tissue and in draining mesenteric lymph nodes (mLN), available at www.gutcellatlas.org. We demonstrate previously unappreciated changes in the proportions and activation status of T and B cells in distinct regions of the healthy human colon from proximal to distal, and relate these differences to the changing microbiota.

## Results

### Microbiome composition differs along distinct colon regions

To create a map of bacterial composition at the mucosal surface of the colon, we performed 16S ribosomal sequencing of swabs from the mucosa surface of the caecum, transverse colon and sigmoid colon of twelve disease-free Caucasian deceased transplant donors (Figure 1a & Supp Table 1). The major gut phyla - *Bacteroidetes, Firmicutes, Proteobacteria* and *Actinobacteria*, were present throughout the gut of each donor (Figure 1b & Supp Table 2). While diversity of OTUs was consistent across the colon, and significant variability existed between donor as previously reported ^15^ (Supp Figure 1a), we did observe changes in the composition of the microbiome. Most notably at the level of phyla, *Bacteroidetes* was more prevalent in sigmoid colon (Figure 1c & Supp Figure 1b). This was mostly attributable to an increase in *Bacteroides*, which dominates the colonic microbiome of individuals on high protein and fat diets typical in Western countries ^16^ (Figure 1c and Supp Figure 1a). Additionally, *Enterococcus* was more prevalent in the proximal colon, and *Coprobacillus* and *Shigella*, typically pathogenic, were more abundant in the distal colon, although these proportions varied considerably between donors (Figure 1b & 1c & Supp Figure 1c).

**Figure 1:**
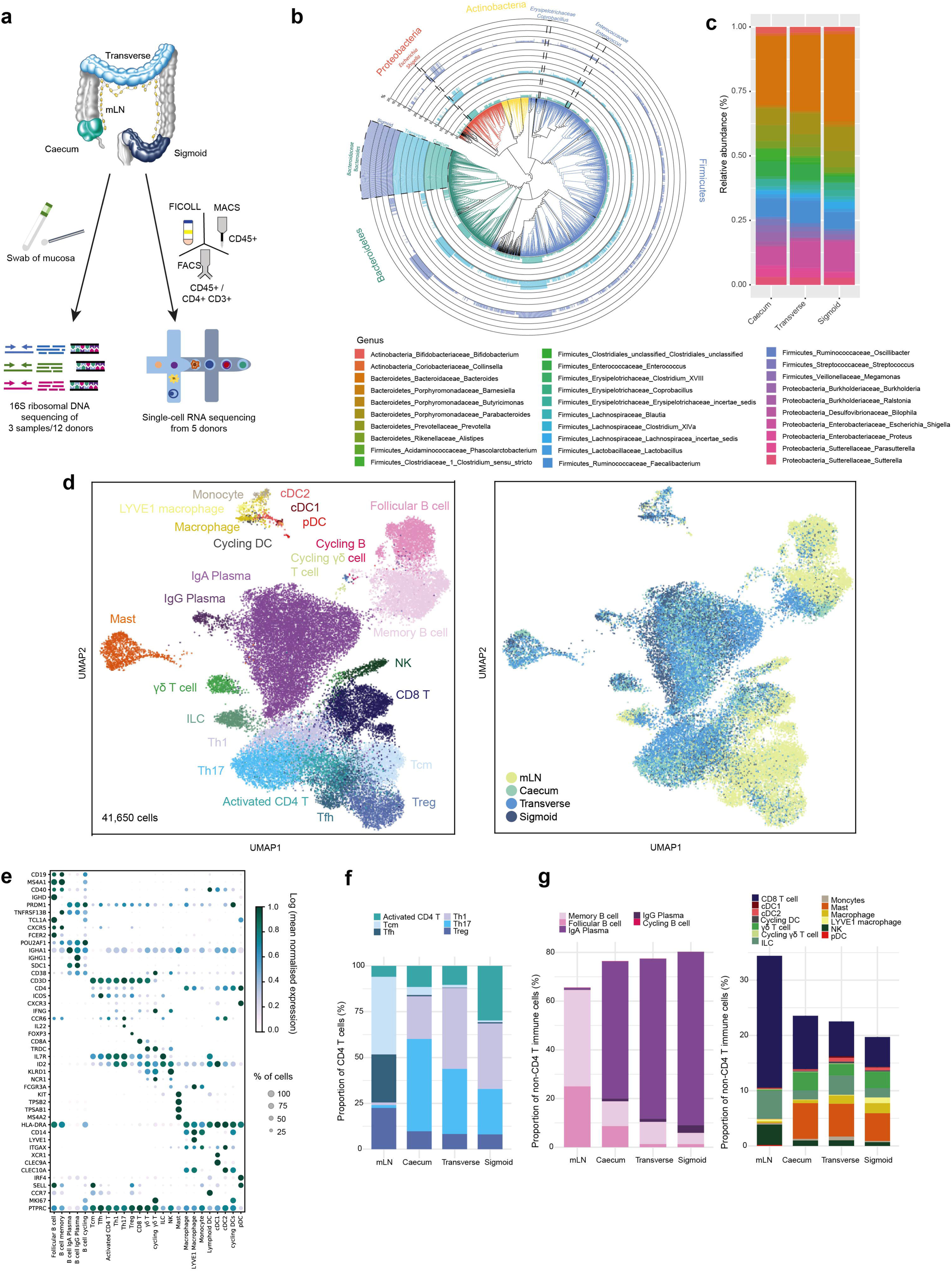
Profiling immune cells along the steady-state colon. a) Workflow for 16S ribosomal sequencing of matching mucosal microbiomes (n=12 donors) and scRNA-seq profiling of immune cells from mesenteric lymph node (mLN), and lamina propria of caecum, transverse colon and sigmoid colon (n=6 donors). b) Phylogenetic tree representing diversity and mean abundance of bacterial species in the caecum, transverse colon and sigmoid colon. Mean abundance was calculated as the percentage of OTU for each species from total as determined by 16S ribosomal sequencing and averaged for twelve donors (black scale). Unassigned OTUs are shown as black branches. Bacteria groups of interest are highlighted. c) Relative abundances of OTUs at genus level of bacteria species in colon as in b. d) Clustering of pooled immune cells visualised in a UMAP plot coloured by tissue of origin (left) and cell type annotation (right). e) Mean expression level and percentage of cells expressing marker genes used to annotate clusters in d. f) Relative percentages of CD4^+^ T subpopulations within all CD4^+^ T cells for each tissue as in d. g) Relative percentages of cell types within all non-CD4^+^ T immune cells for each tissue as in d, with B cells shown in the left panel and all other cell types in the right panel.

Past studies characterising the colonic microbiome typically rely on stool samples, which do not accurately recapitulate the composition of bacteria at the mucosal surface ^17^. Our catalog of mucosal bacteria throughout the colon demonstrates heterogeneity in the microbiome at the mucosal surface from proximal to distal colon, and reveals specific genera with preference for colonising certain colon regions.

### Immune cell heterogeneity in steady-state colon

Next, we sought to determine whether the heterogeneity we observed in the colonic microbiome was accompanied by differences in the adjacent host immune cells. To this end, we generated high-quality transcriptional data from over 41,000 single immune cells from the mLN and lamina propria of caecum, transverse colon and sigmoid colon (Figure 1a). We acquired tissue biopsies from five deceased transplant donors (Supp Table 1). Samples were dissociated to release cells from the mLN and lamina propria of colonic tissue. Immune cells were enriched either by fluorescence-activated cell sorting (FACS) of CD4^+^ T cells (live CD45^+^CD3^+^CD4^+^) and other immune cells (live CD45^+^CD3^-^CD4^-^), or by CD45^+^ magnetic bead selection or Ficoll gradient. Each fraction was then subjected to scRNA-seq (see Methods; Figure 1a & Supp. Figure 1d). Despite the enrichment for immune cells, we captured epithelial cells and fibroblasts and these were computationally removed from further analysis (see Methods).

Pooled analysis and visualisation with Uniform Manifold Approximation and Projection (UMAP) of all four tissues from all donors revealed distinct clusters in the lymphoid and colonic tissues (Figure 1d & Supp Figure 1e). We identified 25 cell types and states in the intestinal lamina propria and mLN (Figure 1d & Supp Table 3 & Supp Table 4), consistent with the immune populations described in recent reports ^18^. Among these were follicular and memory B cells, IgA^+^ and IgG^+^ plasma cells, effector and memory CD4^+^ T cells, T regulatory (Treg) cells, CD8^+^ T cells, γδ T cells, innate lymphoid cells (ILCs), natural killer (NK) cells, mast cells and myeloid cells (Figure 1d). Sub-clustering of myeloid cells showed two distinct populations of conventional dendritic cells: cDC1 expressing *XCR1, CADM1, CLEC9A, BATF3* and *IDO1*, and cDC2 expressing *CLEC10A* and *CD86* (Figure 1d & 1e). In addition, we identified monocytes expressing *CD14* and *CD68*, macrophages expressing *FCGR3A* (gene encoding CD16), LYVE1 macrophages ^19^ and plasmacytoid DCs (pDCs) expressing *IRF4* and *SELL*. B cells, DCs and γδ T cells had *MKI67*^+^ cycling populations suggesting higher rates of proliferation compared to other colonic immune cell populations (Figure 1d & 1e).

To determine how immune cells differ along the colon from proximal to distal regions, we investigated the relative proportions of cell types within mLN and colon regions (Figure 1f & 1g). Since CD4^+^ T cells and all other immune cells were sorted separately during initial tissue processing, our analyses here were also kept separate. As noted from our visual inspection of the UMAP plot (Figure 1d), there were major differences in the cell types present between mLN and the colon. In particular, there were marked differences in the activation and memory status of T and B cells between the colon and lymph nodes, suggesting that these cell types are moulded by their environment (Figure 1f & 1g). In the mLN, CD4^+^ T cells were typically *CXCR5*^+^, *ICOS*^*hi*^ follicular helper cells, and *SELL*^+^ (encodes CD62L), *CCR7*^+^ central memory cells (Figure 1e and 1f). In contrast, colonic CD4^+^ T cells had a more effector phenotype, expressing high levels of the tissue residency marker *CD69* ^*20*^, falling into the Th17 (*CCR6*^+^, *IL22*^+^) or Th1 (*CXCR3*^*+*^, *IFNG*^+^) subtypes. There was an inverted gradient in the relative proportion of Th17 and Th1 cells, with the caecum dominated by Th17 cells that reduced in frequency in the transverse colon, and still further in the sigmoid colon, and Th1 cells following the opposite trend, being more abundant in the sigmoid colon (Figure 1f). This distinct distribution of colonic Th1 and Th17 cells is concordant with spatial variation of the microbiome, hinting at a relationship between the two populations.

B cells in the mLN were predominantly *CD19*^*+*^, *MS4A1*^*+*^ (encodes CD20), *CD40*^+^, *TNFRSF13B*^+^ (encodes TACI), *CD38*^*-*^ memory or *CXCR5*^+^, *TCL1A*^+^, *FCER2*^+^ (encodes CD23) follicular B cells (Figure 1e and 1g). In contrast, in the three regions of the colon, the main population of B cells were *SDC1*^+^ (encodes CD138), *CD38*^+^, plasma cells. Curiously, plasma cells were enriched in the sigmoid colon relative to both the caecum and transverse colon, whilst the proportion of memory B cells was lower in the sigmoid colon (Figure 1g). This suggests that conditions in the sigmoid colon may favour the generation of plasma cells rather than memory B cells from germinal centre responses in this region, or that the tissue contains more plasma cell niches.

### T helper cells disseminate through the colon and adopt region-specific transcriptional profiles

We next investigated CD4^+^ T effector cells across the colon. Correlation analysis of Th1 and Th17 cells between different colon tissues revealed high transcriptional similarity between these effector cell subtypes (Figure 2a), with mLN versus peripheral tissue signature accounting for the greatest amount of variability (Spearman’s corr = 0.88). Within the effector T cells of the colon, transverse colon and caecum cells cluster by T helper subtype. Surprisingly, Th1 and Th17 cells of the sigmoid colon did not cluster with their respective effector subtypes from other regions. Differential gene expression analysis between the effector cells in the sigmoid colon versus those in caecum and transverse colon revealed higher expression of activation-related molecules including *TANK* (TRAF family member associated NF-kB activator) (adjusted P < 10^−10^), *CD83* (adjusted P < 10^−10^) and *PIM3* (adjusted P < 10^−8^) (Figure 2b). Expression of *CCL20*, encoding the ligand for CCR6 that is expressed by epithelial and myeloid cells more highly in small intestines than colon ^21^, was also slightly increased by T helper cells of the proximal colon (Figure 2b), although this is likely due to higher abundance of *CCL20*^+^ Th17 cells at this site (Figure 1f & Supp Figure 2a). Conversely, sigmoid colon effector T cells showed higher expression of *KLF2* (adjusted P < 10^−14^) (Figure 2b) that encodes a transcriptional factor that transactivates the promoter for Sphingosine-1-phosphate receptor 1 (S1PR1) and is critical for T cell recirculation through peripheral lymphoid tissue ^22^, *LMNA* (adjusted P < 10^−47^) that is reported to promote Th1 differentiation ^23^ and *EEF1G* (adjusted P < 10^−13^), a driver of protein synthesis ^24^.

**Figure 2:**
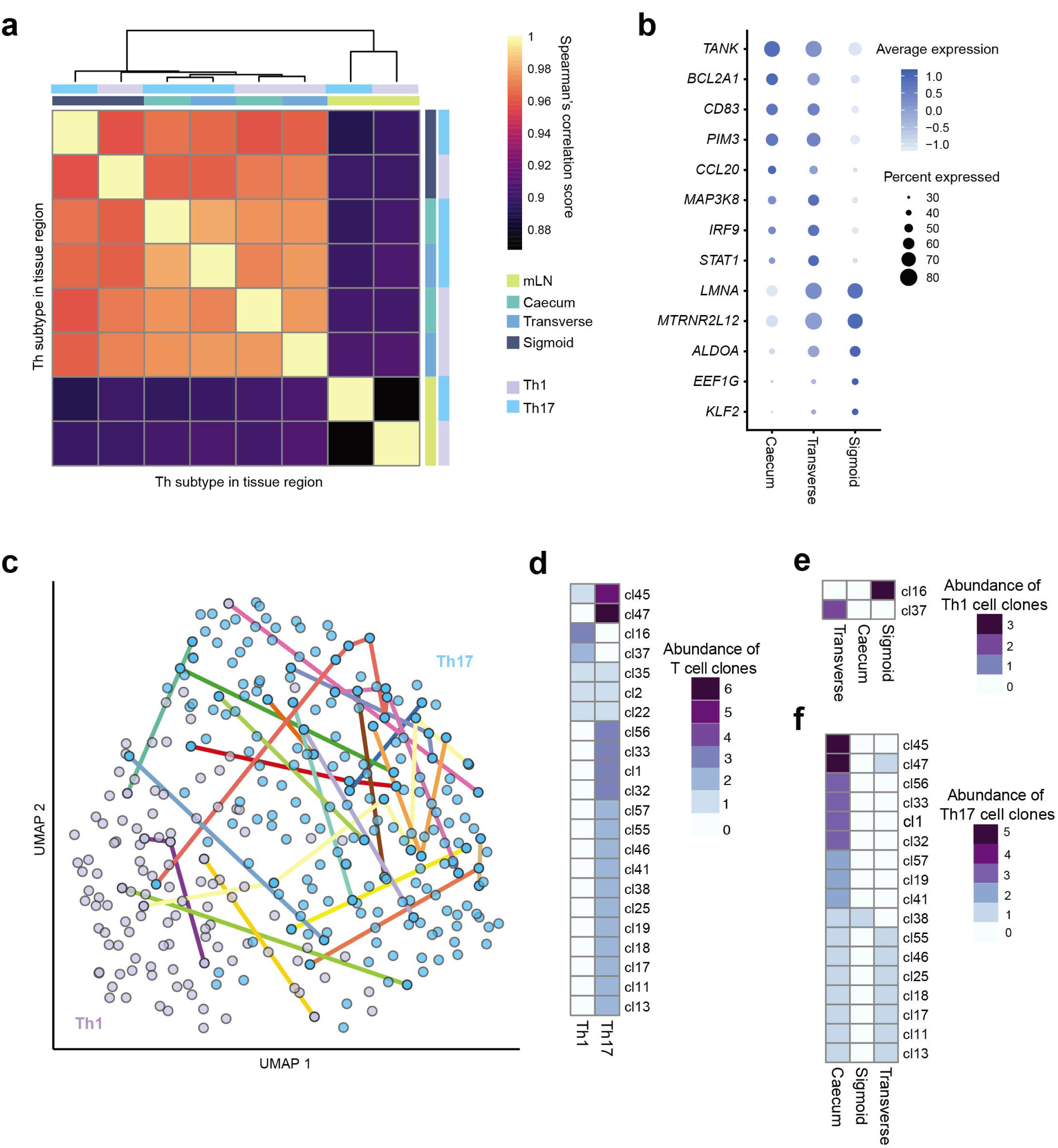
Dissemination of T helper cells in colon and region-determined transcriptional profiles. a) Correlation matrix of mean transcriptional profiles of Th1 and Th17 cells from caecum, transverse colon, sigmoid colon and mLN (n=5 donors). b) Mean expression level of differentially expressed genes of T helper cells between caecum, transverse colon and sigmoid colon. c) UMAP projection of Smartseq2 profiled Th1 and Th17 cells of the caecum, transverse colon and sigmoid colon (n=1 donor). Colour lines connect cells sharing the same CDR3 sequence. d) Heatmap of numbers of members within clonal families in Th1/Th17 subsets (left) and colon region (right).

Next, we looked into the clonal relationships between T helper cells. We performed plate-based Smartseq2 on TCRα/β^+^ FACS-sorted cells from colon regions and mLN of a sixth donor to capture paired gene expression and TCR sequences from individual T cells. Clonal groups were shared between Th1 and Th17 subtypes, supporting the notion that effector fate of CD4^+^ T cells is determined after their initial activation ^25^. Additionally, clonal expansion was observed by Th1 cells of the sigmoid colon (Figure 2C), in line with greater abundance in this tissue. Likewise, clonal expansion of Th17 cells was greatest in the caecum matching accumulation seen with the droplet-based scRNA-seq analysis. Several Th1 and Th17 clonal sisters were shared between clonal sites (Figure 2C), evidence that Th clones disseminate to distant regions of the colon.

Together these data demonstrate region-specific transcriptional differences relating to activation and tissue migration in Th1 and Th17 cells of the proximal and sigmoid colon. Identification of clonal sharing between these colon regions supports the idea that these observed transcriptional differences are due to cell-extrinsic rather than intrinsic factors.

### Activation trajectory of colonic CD4^+^ T regulatory cells

Treg cells are known to play a role in balancing the immune activity of other CD4 T cell subsets. Documented Treg cell activation by *Clostridium spp*.^10^ and previous descriptions of tissue-specific transcriptional profiles ^26^ inspired us to interrogate these cells in greater detail. Firstly, we noted that the relative proportion of Treg cells did not change significantly from proximal to distal colon (Figure 1f). We then investigated whether the transcriptional profiles of Tregs from different compartments indicated distinct activation states or functionalities.

As we previously observed in the mouse ^26^, sub-clustering of Treg cells from the mLN revealed major populations of central Tregs and effector Tregs (Figure 3a)^26^. Central Tregs were defined by highest expression of *SELL*, while effector Tregs were characterised by genes associated with the TNFRSF-NF-κB pathway (*TNFRSF9* and *TNF*) (Supp Figure 3a). In addition, we observed a population, previously termed non-lymphoid tissue-like Tregs (NLT-like Treg). These have characteristics of non-lymphoid tissue Treg cells, including high expression of *FOXP3, PRDM1, RORA, IL2RG, IL2RA* and *CTLA4* (Figure 3a & Supp Figure 3a). A fourth population, termed Treg-like cells, lacked the conventional markers of Treg cells - *FOXP3* and *IL2RA* - but clustered more closely with Treg cells than conventional T cells (Figure 3a and Supp Figure 3a). These expressed the highest levels of *PDCD1* (gene encoding PD-1) among Treg populations and, uniquely, the transmembrane protein *MS4A6A* (Supp Figure 3a). This population of Treg-like cells could represent a Treg population that transiently loses FOXP3 expression ^27^.

**Figure 3:**
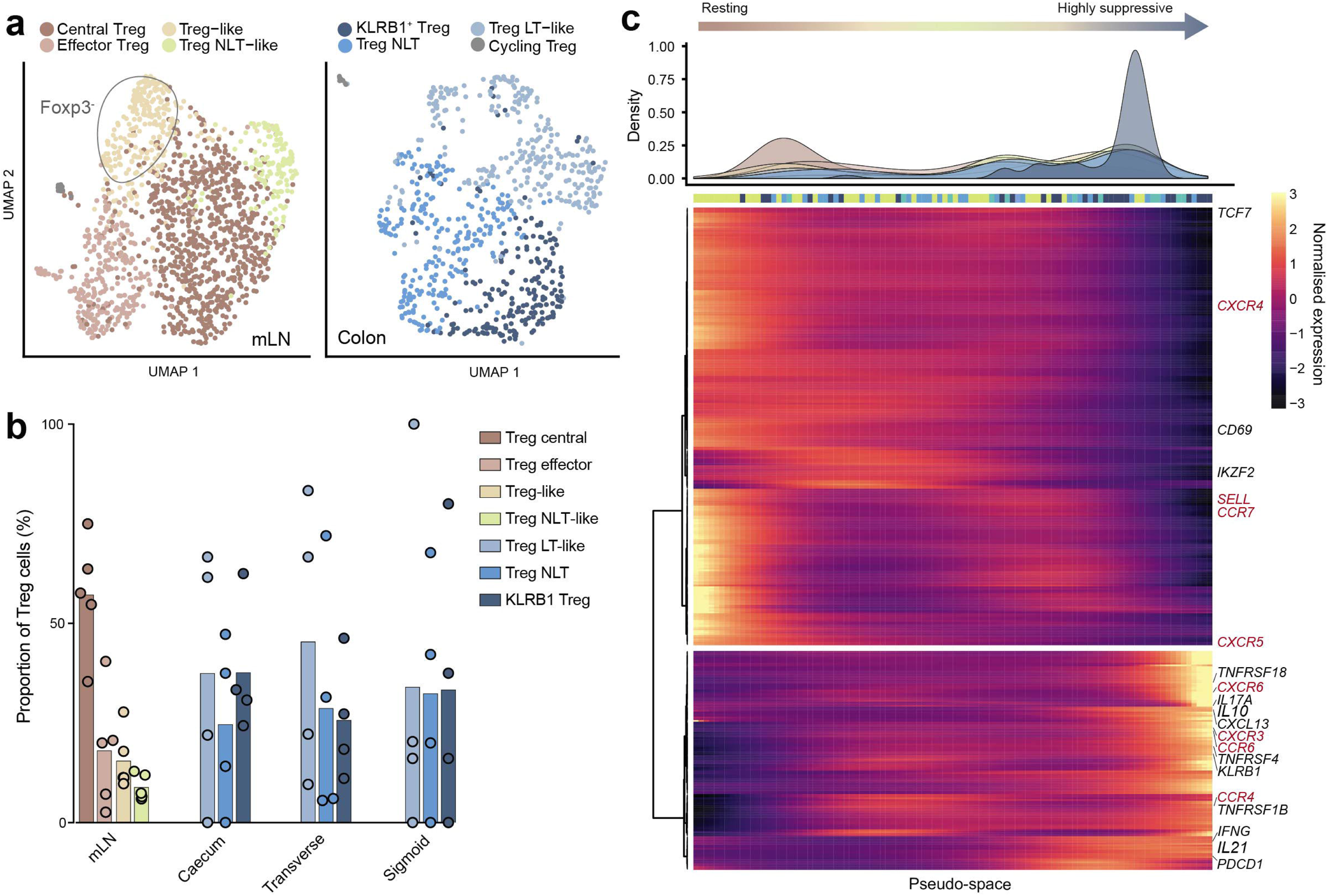
Treg activation pathway from lymphoid to peripheral tissue. a) UMAP visualisation of Treg subtypes in mLN (left) and colonic regions (right) (n=5 donors). b) Relative proportions of Treg subsets within all Treg cells from mLN and colon tissue regions. Bars show mean proportion across all donors (circles). c) Density of Treg subclusters as in a across ‘pseudospace’ (top) and expression kinetics of genes contributing to pseudospace smoothed into 100 bins (filtered by qval < 0.001 and expression in >15 cells). Top bar shows most represented tissue within each bin. Various dynamically expressed immune-related molecules are annotated, with key genes coloured red.

In the colon, Tregs clustered into three populations as previously described in mouse (Figure 3a) ^26^. KLRB1^+^ (also known as CD161) Treg cells were characterised by expression of *LAG3, IL2RA, CTLA4, KLRB1, ICOS* and *FOXP3* (Supp Figure 3a) suggesting a robust regulatory function, analogous to the peripherally-derived induced Treg cells described in mouse ^28^ and similar to CD161^+^ Tregs described in human colon tissue ^20,29^. Non-lymphoid tissue Treg (NLT Treg) cells express *IKZF2, GATA3* and *DUSP4* (Supp Figure 3a) consistent with the profile of thymic-derived Tregs ^30^. The third population exhibited a profile reminiscent of lymphoid tissue Tregs with expression of *SELL, CCR7, TCR7, CXCR5* and *RGS2*, and were termed Lymphoid Tissue-like Treg (LT-like Treg) cells (Supp Figure 3a). LT-like Treg cells are likely newly arrived in the colon tissue from mLN. A small number of *Ki67*^*+*^ Treg cells were also identified in the colon and mLN (Figure 3a & Supp Table 3). The proportions of these subsets varied by donor, but were mostly consistent between colon regions (Figure 3b).

The presence of NLT-like Treg cells in the mLN and LT-like Treg cells in the colon, both with profiles suggesting migration, led us to recreate this migration pathway *in silico*. We ordered all Tregs along “pseudo-space” using Monocle2. This gave rise to a smooth pseudo-space trajectory from resting central Tregs in the mLN to highly suppressive Tregs in the colon (Figure 3c). As seen in the mouse ^26^, NLT-like Tregs and LT-like Tregs blended in the middle of the trajectory, in accordance with these cells representing transitioning and migratory populations between lymphoid and peripheral tissues. In order to understand which gene signatures drive the migration and tissue adaptation of Tregs in human tissues, we determined the genes changing along the previously calculated ‘pseudo-space’ (Figure 3c). Genes expressed at the beginning of pseudo-space included *SELL, CCR7* and *CXCR4*, permitting entry into lymph nodes (Figure 3c). At the end of pseudo-space, the most highly expressed genes were *FOXP3, IL2RA, CTLA4, IL10* and *LAG3.* These largely suppressive genes were co-expressed with TNF receptors (*TNFRSF4, TNFRSF14, TNFRSF18, TNFRSF1B*) indicating a reliance on the TNFRSF-NF-κB axis. Chemokine receptors *CXCR3, CXCR6, CCR6* and *CCR4* were also expressed by Tregs cells in the periphery, matching previous reports of Th1/Th17-like Tregs cells in the colon (Figure 3c)^26^.

Together, these results highlight heterogeneity in Treg cell states in mLN and colon, and reveal a possible Foxp3-transiently absent population. We also infer a continuous activation trajectory of these Treg cell states between draining lymph nodes and colon, and highlight genes regulating Treg cell migration between tissues and their adoption of Th-like profiles.

### B cells display a proximal-to-distal activation gradient

Following from our observations in CD4^+^ T cells across the colon, we next focused on humoral responses by performing a more in-depth analysis of B cells in different colon regions. We compared transcriptional profiles of plasma cells between different colonic regions. This analysis revealed *CCL3* and *CCL4* as highly enriched in caecal plasma cells (log fold change of 0.61 and 0.70 and adjusted P <10^−28^ and <10^−9^ respectively) (Figure 4a). These chemokines are secreted by B cells in response to BCR activation ^31^ and result in the migration of CCR5-expressing cells such as T cells ^32,33^ and monocytes ^34^ to the tissue microenvironment. This suggests that BCR cross-linking and signalling may be more prominent in the proximal colon. Caecal plasma B cells were also enriched for *CXCR4* (log fold change of 0.44, adjusted P < 10^−10^) (Figure 4a), a chemokine receptor highly expressed by germinal centre (GC) B cells and important for the movement of plasmablasts to the GC-T zone interface post-GC responses ^35^.

**Figure 4:**
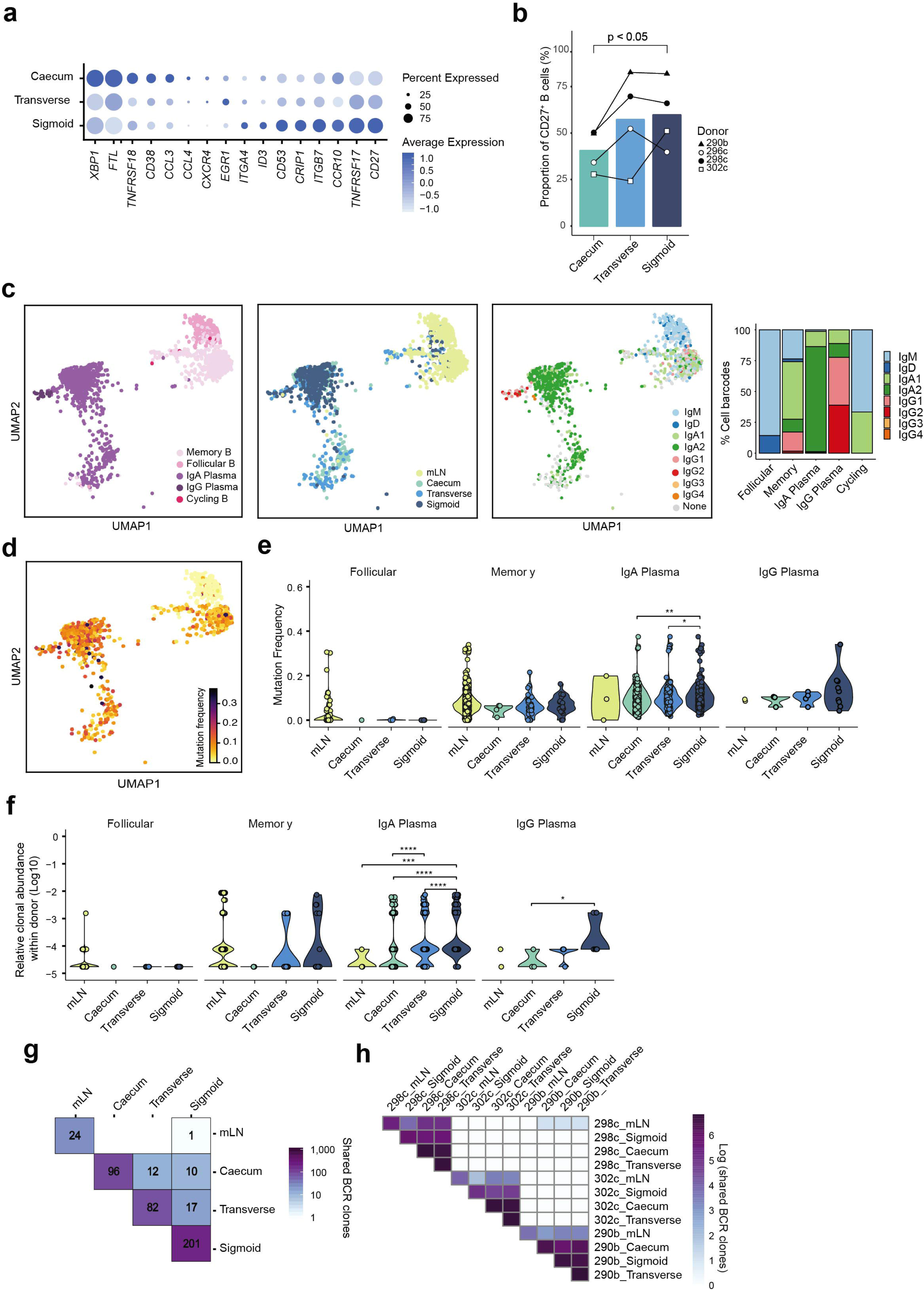
B cells are more abundant, clonal expanded and mutated in the sigmoid colon. a) Log normalised mean transcript counts of key differentially expressed activation and tissue-migration genes by IgA^+^ plasma cells in caecum, transverse colon and sigmoid colon (n=5 donors). b) Proportion of CD27^+^ B cells of total B cells from colon regions determined by flow cytometry (n=4 donors). Bar represents mean proportion and connected points represents values of each donor. Analysis is a two-tailed paired t test. c) Clustering of B cells for which matched single-cell VDJ libraries were derived using 10x Genomics 5’ scRNA-seq (n=2 donors). B cells are visualised by UMAP projection coloured by cell type annotation (left), tissue of origin (middle) and antibody isotype (right). Antibody isotype frequencies per annotated cell type across both donors are depicted as a bar plot (far right). d) Somatic hypermutation frequencies of single-cell VDJ-derived IgH sequences as in c. e) Quantitation of somatic hypermutation frequencies of single-cell VDJ-derived IgH sequences from B cell types and gut regions. f) Estimated clonal abundances per donor for members of expanded B cell clones in B cell types and gut regions. g) Co-occurrence of expanded B cell clones identified by single-cell VDJ analysis shared across colon regions as in c. Numbers reflect binary detection event rather than the number of members per clone shared. h) Co-occurrence of expanded B cell clones identified by bulk B cell receptor (BCR) sequencing shared across mLN, caecum, transverse colon and sigmoid colon (n=3 donors). Rows and columns in g and h are ordered by hierarchical clustering.

Amongst the genes more highly expressed by plasma cells in the sigmoid colon was *CD27* (Figure 4a; log fold change of 0.24; adjusted P <10^−40^), a member of the TNF receptor family that is expressed by memory B cells and even more highly by plasma cells ^36^. We confirmed differential expression of CD27 at the protein level and additionally observed a proximal-to-distal gradient of increasing CD27 expression by plasma cells in the colon (Figure 4b). Targeted homing of B cells from their site of activation in lymphoid tissues to the colonic lamina propria relies on signalling through CCR10 and its cognate ligand, CCL28, and integrin α*4*β*7* ^37,38,39^. *CCR10* (log fold change of 0.24; adjusted P < 10^−16^), *ITGA4* (log fold change of 0.58; adjusted P < 10^−24^) and *ITGB4* (log fold change 0.57; adjusted P < 10^−108^) were also more highly expressed by sigmoid colon IgA^+^ plasma cells (Figure 4a).

To determine whether the B cell clonal repertoire changes across the colon, we took advantage of the paired single-cell VDJ-sequencing data available from two donors for which scRNA-seq libraries were generated using 10x Genomics 5’ chemistry. We confirmed the expression of IgM and IgD isotypes by follicular B cells, IgG1 and IgG2 by IgG^+^ plasma cells, IgA1 and IgM expression by memory cells in the mLN and predominantly IgA2 expression by plasma cells of the colon (Figure 4c). The mutation frequency of the heavy chain variable region was greatest in the plasma cells followed by memory B cells, indicating more somatic hypermutation by these cell types compared to the naive follicular B cells (Figure 4d). Additionally, while mutational frequency was consistent across colon regions and mLN for memory and follicular B cells, it was significantly increased in IgA^+^ plasma cell of the sigmoid colon compared to the other colon regions (Figure 4e). IgG^+^ plasma cells also showed a trend towards increased mutational frequency in the sigmoid colon, however their numbers were limiting (Figure 4e).

We then identified clonally-related B cells and plasma cells examine clonal expansion dynamics of different cellular populations throughout the gut. Clonal expansion was evident in memory B cells, IgA^+^ plasma cells and IgG^+^ plasma cells (Figure 4f). While the relative abundance of clonal groups did not differ across the colon regions of memory B cells and IgG^+^ plasma cells, again this was greatest for IgA^+^ plasma cells in the sigmoid colon (Figure 4f). This was supported by bootstrapped VDJ sequence diversity analysis of clonally-related IgA^+^ plasma cells, which showed that the diversity of BCR sequences was consistent between donors and that there was a trend for decreased diversity (consistent with higher rates of clonal expansion) of IgA^+^ plasma cells in the sigmoid colon (Supp Figure 4a & 4b). While some clones were shared between B cell types (i.e. memory and IgA^+^ plasma), indicating that alternate B cell fates can derive from a single precursor cell, most expanded clones within the gut were of the same cell type (Supp Figure 4c).

Finally, we found many examples of B cell clones shared between all three colonic regions, and to a lesser extent the mLN, for both donors (Figure 4g), indicating dissemination of B cells throughout the colon as previously reported ^40^. Our observation of B cell dissemination throughout the colon was replicated with bulk BCR sequencing from whole tissue (Figure 4h) ^41^. Interestingly, increased clonal variability of sigmoid colon B cells was reflected in a greater spread of BCR variable chains expressed compared with caecum and transverse colon (Supp. Figure 4d), altogether suggesting a more active response in the distal versus proximal colon.

These data indicate a highly activated state of plasma cells in the distal colon compared with proximal colon plasma cells, characterised by greater accumulation, somatic hypermutation, clonal expansion and stronger homing to the colon mucosa.

### IgA is more responsive to specific bacterial species in the sigmoid colon

Reports by Macpherson et al. have shown that IgA is secreted by plasma cells in response to the presence of specific bacteria species, rather than as a general response to the microbiome ^42^. In light of this, we wondered whether the increased plasma cell activation we observed in the sigmoid colon was linked to the differences in reactive bacteria species. To this end, we assessed IgA-opsonisation of bacteria from the donor microbiome samples (Figure 5a). Surprisingly, a greater proportion of bacteria from the sigmoid colon was positive for IgA-binding compared with bacteria of the caecum and transverse colon (Figure 5b). Furthermore, shotgun sequencing of IgA-opsonised bacteria revealed a richer community of species in the sigmoid colon (Figure 5c & Supp Table 5). Interestingly, diversity of IgA-bound bacteria, which considers relative abundance of each species, was lower in the sigmoid colon compared to caecum (Figure 5d).

**Figure 5:**
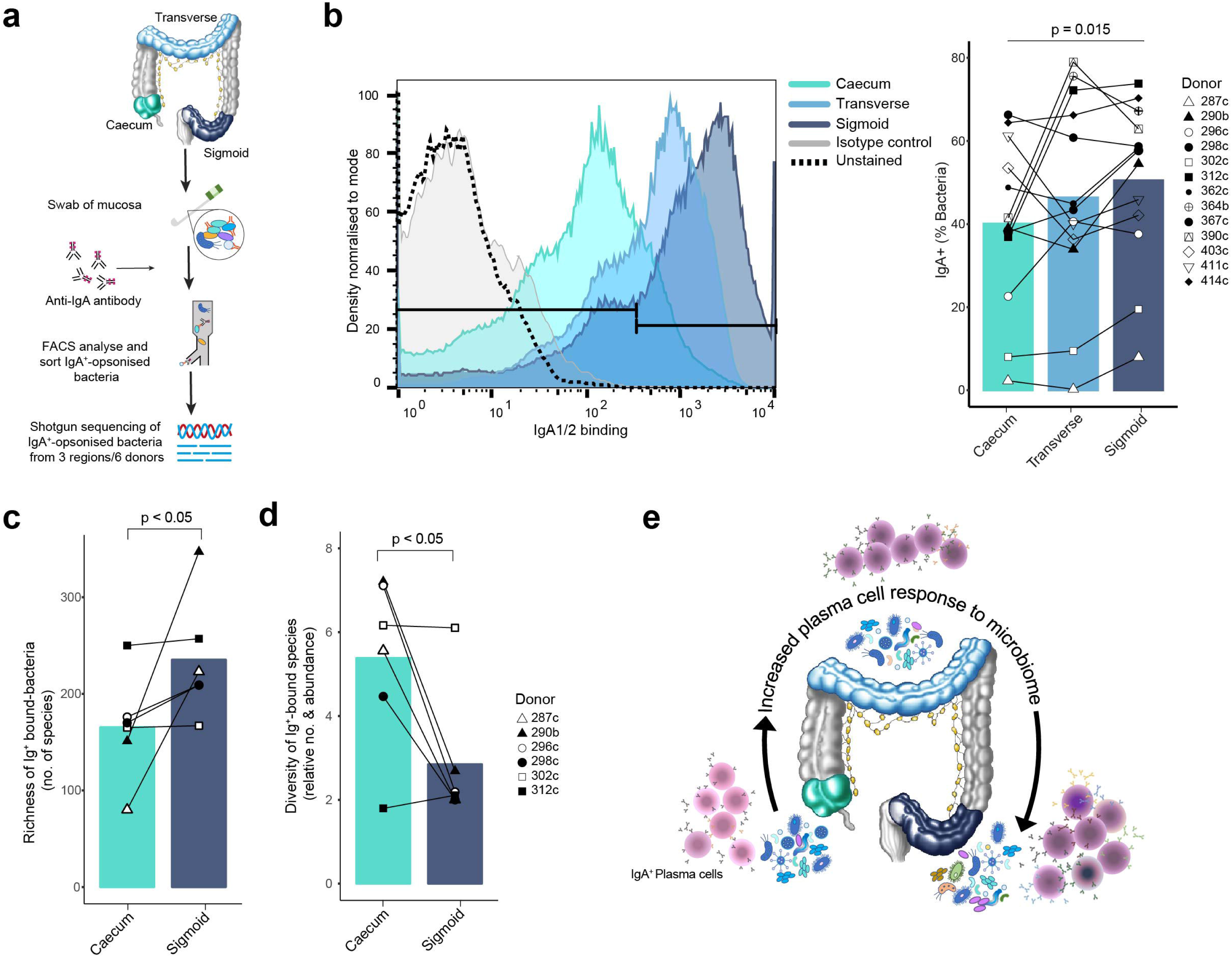
Increasing number of microbiome species recognised by antibodies in the sigmoid colon. a) Experimental workflow for assessing Ig-opsonised colon bacterial species. b) Representative histogram of IgA1/2-bound Hoechst^+^ bacteria and summary plot of bound bacteria as a proportion of total bacteria (n=13 donors). Positive binding is set against an isotype control. c) Richness of bacteria species determined as the number of unique species and d) diversity of species identified from shotgun sequencing of Ig-opsonised bacteria from b (n=6 donors). e) Summary schematic showing proximal-to-distal increasing in accumulation, targeted migration, clonal expansion and mutation frequency of IgA^+^ plasma cell, and associated increasing diversity of reactive bacteria. P values were calculated using two-tailed paired ANOVA with multiple comparisons (b) and one-tailed paired ANOVA with multiple comparisons (c and d).

These data suggest that, compared with the caecum, IgA^+^ plasma cells of the sigmoid colon respond to a rich and unevenly represented community of bacterial species, likely contributing to their increased activation status, strong homing to the colon and trend towards greater clonal diversity (Figure 5e).

## Discussion

In this study we performed the first simultaneous assessment of the colonic mucosal microbiome and immune cells in human donors at steady-state. This enabled us to compare lymphoid and peripheral tissue immunity and explore how immune cells and their neighbouring microbiome change along the colon within the same individuals. In doing so, we highlight previously unappreciated regional differences in both cellular communities. Our unique annotated colon immune single-cell dataset is available at www.gutcellatlas.org, where users can visualise their genes of interest.

We describe a shift in the balance of T helper subsets, with a predominance of Th17 in the caecum and Th1 in the sigmoid colon. Decreasing abundance of Th17 cells has similarly been shown from proximal small intestine to colon of mice ^7^. Additionally, simultaneous increase of the genus *Bacteroides* and Th1 numbers in the sigmoid colon are in line with findings that polysaccharide from *Bacteroides fragilis* preferentially induces Th1 differentiation in the intestine of germ-free mice ^11^. An alternative or complementary explanation for skewed T helper proportions and variation in transcriptional profiles is offered in a study by Harbour et al., which demonstrated that Th17 cells can give rise to a IFNy^+^ ‘Th1-like’ cell in response to IL-23 production by innate cells ^43^. These findings demonstrate the complexity of external signals shaping colonic Th responses leading to regional changes in their numbers and differentiation.

In contrast to conventional T cells, Treg cells are evenly represented across the colon. Treg cell subpopulations within the mLN and colon tissue are analogous to those we have recently described in mouse ^26^. We also identify an additional population, termed Treg-like cells, reminiscent of a CD25^-^FOXP3^lo^PD1^hi^ Treg population in the peripheral blood, although the latter was described to also express Ki67 ^44^. This population could represent uncommitted Treg cells experiencing transient loss of FOXP3 while retaining regulatory potential ^27^ or permanent loss FOXP3 and adoption of a more pro-inflammatory phenotype after repeated stimulation ^45^. Our pseudo-space analysis of Treg cells suggests a continuum of activation states from resting cells in the lymphoid tissue through to highly suppressive cells in the periphery. We identify genes underlying this transition including chemokine receptors that are strongly expressed on arrival in the intestine that enable interaction with, and suppression of, Th1 and Th17 cells ^46^. Similarly to mouse, pseudospace from lymphoid to peripheral tissue correlates with transcriptional markers of migratory potential and suppression of effector cells.

We find that IgA^+^ plasma cells are more abundant and have greater expression of colon-specific migration markers (*CCR10, ITGA4* and *ITGB7*) in the sigmoid colon compared to the proximal colon. This adds finer resolution to previous reports describing increasing abundance of IgA^+^ plasma cells from the small intestine to the colon ^47^. Our B cell repertoire analysis demonstrates extensive clonal expansion within each colon region and to a lesser extent between regions, arguing for colonic dissemination of B cells from the same precursor pool, followed by local expansion. Furthermore, more clonal sharing existed between regions of the colon than with mLN, consistent with recent work showing that while mLN clones can also be detected in blood, the intestines are host to a unique B cell clonal network ^48^. Sigmoid colon plasma cells, in particular, exhibit greater mutational abundance and clonal expansion. Previous work has shown the mutational frequency of B cells is consistently high between duodenum and colon and are primarily driven by dietary and microbiome antigens respectively ^49^. Thus, we suggest that enhanced plasma cell accumulation, mutation and expansion in the sigmoid colon is in response to continued stimulation from the local microbiome. This may happen through increased engagement of sigmoid colon plasma cells in T cell-mediated germinal centre reactions in gut-associated lymphoid structures or in T cell-independent somatic hypermutation in local isolated lymphoid follicles ^50^ followed by local expansion. Yet the exact mechanisms require further study.

Finally, we show greater IgA binding to the microbiome in the sigmoid colon compared to proximal sites (Figure 5b). One possible explanation for this observation is the accumulation of upstream secreted IgA and IgA-bound bacteria in the sigmoid colon. However, our simultaneous observation of enhanced plasma cell responses in the sigmoid colon suggests that IgA is locally produced as a result of immune poising. Possible scenarios contributing to a richer immune-reactive microbiome in the sigmoid colon are bacteria derived externally via the rectum. Alternatively, environmental pressures (i.e. lower water and nutrient levels ^4^) could restrict outgrowth of dominant gastrointestinal species of the proximal colon, providing space for smaller communities of opportunistic species. The IgA response in the colon is antigen-specific rather than a general response to the presence of bacteria ^51,42^. Thus, the overall increased number of unique species recognised by host IgA antibodies in the sigmoid colon is fitting with the enhanced clonal expansion and mutation of plasma cells at this site.

Together, our simultaneous analyses of microbiome and neighbouring immune cells highlight the significance of environmental signals in shaping and maintaining regional adaptive immune cell composition and function in the intestine at steady-state. Dysregulation of T helper cells ^52^ and plasma cells ^53,54,55^ has been implicated in susceptibility to inflammatory bowel disease. Observations of the linked compartmentalisation of these immune cells and microbial species along the colon at steady-state may provide a platform for understanding the mechanisms underpinning the tropism of different intestinal diseases to specific regions of the gut, such as Crohn’s disease and ulcerative colitis.

## Acknowledgements

The authors gratefully acknowledge the support received from the Sanger Single Flow Cytometry Facility, Sanger Cellular Genetics Department IT team and Core Sanger Sequencing pipeline. We thank Dr Foad Rouhani and Tomas Castro-Dopico for insightful discussions about project design and experimental design and Dr Krishnaa Mahbubani for help with collection of human tissue. We acknowledge Jana Eliasova for graphical images. This research was supported by funding from Wellcome (grant no. WT206194) and European Research Council (grant no. 646794). K. R. J. receives financial support from Christ’s College, University of Cambridge. T. G. was funded by the European Union’s H2020 research and innovation programme “ENLIGHT-TEN” under the Marie Sklodowska-Curie grant agreement 675395. H. W. K. was funded by a Sir Henry Wellcome PostDoctoral Fellowship (grant no. 213555/Z/18/Z). B. R. B. was funded by the NIHR Cambridge Biomedical Research Centre. We are grateful to the deceased organ donors, donor families and the Cambridge Biorepository for Translational Medicine for access to the tissue samples. This publication is part of the Human Cell Atlas - www.humancellatlas.org/publications.

## Author contributions

K. R. J. initiated this project, designed and performed scRNA-seq and microbiome experiments, analysed data and wrote the manuscript. T. G. analysed bulk BCR-seq data and contributed extensively to scRNA-seq data analysis. R. E. contributed to data interpretation and data analysis. N. K. and E. L. G. analysed 16S ribosomal sequencing and metagenomics data. M. D. S assisted in microbiome related experiments. H. W. K. and L. K. J. analysed the 10x Genomics VDJ datasets and contributed to generation of figures. B. R. B. and K. S. P. carried out tissue collection. J. R. F. designed FACS sorting panel and assisted in FACS sorting. V. P. assisted in bulk BCR library preparation and analysis. L. B. J., O. S., S. H. and J. L. J. dissociated tissues from donor 390c. K. S. P. carried out scRNA-seq read alignment and quality control. S. C. F., K. B. M. and M. R. C. designed experiments and interpreted data. T. D. L. and S. A. T. initiated and supervised the project and interpreted data. All authors edited the paper.

## Methods

### Colon and mesenteric lymph node tissue retrieval

Human tissue was obtained from deceased transplant organ donors after ethical approval (reference 15/EE/0152, East of England Cambridge South Research Ethics Committee) and informed consent from the donor family. Fresh mucosal tissue from the caecum, transverse colon and sigmoid colon, and lymph nodes from the intestine mesentery, were excised within 60 minutes of circulatory arrest and colon tissue preserved in University of Wisconsin (UW) organ preservation solution (Belzer UW® Cold Storage Solution, Bridge to Life, USA) and mLN stored in saline at 4°C until processing. Tissue dissociation was conducted within 2 hours of tissue retrieval. Four individuals (287c, 296b, 403c and 411c) had received antibiotics in the two weeks prior to death (Supp. Table 1).

### Tissue dissociation for FACS separation and MAC separation

Tissue pieces from donors 290b, 298c, 302c, 364b and 411c were manually diced and transferred into 5mM EDTA (Thermo Fisher Scientific)/1mM DTT/10mM HEPES (Thermo Fisher Scientific)/2% FBS in RPMI and incubated in a shaker (∼200rpm) for 20 minutes at 37°C. Samples were briefly vortexed before the media renewed and incubation repeated. Tissue pieces were washed with 10mM HEPES in PBS and transferred into 0.42mg/ml Liberase DT (Roche)/0.125 KU DNase1 (Sigma)/10mM HEPES in RPMI and incubation for 30 minutes at 37°C. The digested samples were passed through a 40um strainer and washed through with FBS/PBS.

### Fluorescence-activated cell sorting

Cells from donor 290b, 298c and 302c were pelleted and resuspended in 40% Percoll (GE Healthcare). This was underlayed with 80% Percoll and centrifuged at 600g for 20 minutes with minimal acceleration and break. Cells at the interface were collected and washed with PBS. Cells were stained for fluorescent cytometry using Zombie Aqua Fixable Viability Dye (Biolegend), CD45-BV605 (clone HI30; Biolegend), CD3-FITC (clone OKT3; Biolegend), CD4-BV421 (clone SK3; Biolegend), CD8-PE-Cy7 (clone SK1; Biolegend), CD19-APC-Cy7 (clone HIB19; Biolegend), IgD-PE Dazzle (clone IA6-2; Biolegend), CD27-BV711 (clone M-T271; Biolegend), HLA-DR-BV785 (clone L243; Biolegend), CD14-APC (clone 63D3; Biolegend) and CD11c-PE (clone 3.9; eBioscience). Non-CD4^+^ T immune cells were sorted as live singlet CD45^+^, CD3- and CD4^-^. CD4^+^ T cells were sorted as live singlet CD45^+^, CD3^+^ and CD4^+^. Each faction was manually counted using 0.4% Trypan Blue (gibco) and a haemocytometer and diluted to 500 cells/ul in PBS. Sorting was carried out on a BD FACS ARIA Fusion. Analysis of FACS data was conducted with FlowJo Software package.

### MACS cell enrichment

Cells from donor 411c were pelleted for 5mins at 300g and resuspended in 80ul of ice-cold MACS buffer (0.5% BSA (Sigma-Aldrich Co. Ltd), 2mM EDTA (ThermoFisher) in DPBS (Gibco)) and 20ul of CD45 Microbeads (Miltenyi Biotech). Cells were incubated for 15 minutes at 4°C before being washed with 2ml of MACS buffer and centrifuged as above. Cells were resuspended in 500ul of MACS buffer and passed through a pre-wetted MS column on QuadroMACS magnetic cell separator (Miltenyi). The column was washed 4 times with 500ul of MACS buffer, allowing the full volume of each wash to pass through the column before the next wash. The column was removed from the magnet and the cells eluted with force with 1ml of MACS buffer into a 15ml tube. Cells were pelleted as above and cell number and viability were determined using a NucleoCounter NC-200 and Via1-Cassette (ChemoMetec). Cells were resuspended at 500 cells/ul in 0.04% BSA in PBS.

### Tissue dissociation for Ficoll separation

Tissues from donor 390c were manually diced and <5.0g was added per Miltenyi C tube with 5mL tissue dissociation media (1x Liberase TL (0.13U/mL/DNase; Roche) and Benzonase nuclease (10U/mL; Merck) in 1% FCS and 20mM HEPES in PBS (Lonza). Samples were dissociated with GentleMACS Octo for 30 minutes homogenising/37°C cycle. Enzymatic digestion was stopped with the addition of 2mM EDTA in tissue dissociation media. Digested samples were then passed through a 70uM smart strainer (Miltenyi) before being washed with PBS and pelleted at 500g for 10 minutes. Cells were resuspended in PBS, layered onto FicollPaque Plus (GE Healthcare) and spun at RT 400g for 25 minutes. Mononuclear cells were retrieved from Ficoll layer and washed with PBS. Cells were filtered through 0.2uM filter (FLowmi cell strainers, BelArt). Cells were manually counted using a haemocytometer and diluted to a concentration of 1000 cells/*µ*l in 0.04% BSA in PBS.

### 10x Genomics Chromium GEX library preparation and sequencing

Cells were loaded according to the manufacturer’s protocol for Chromium single cell 3’ kit (version 2) or 5’ gene expression (version 2) in order to attain between 2000-5000 cells/well. Library preparation was carried out according to the manufacturer’s protocol. For samples from donors 290b, 298c and 302c, eight 10x Genomics Chromium 3’ libraries were pooled sequenced on eight lanes of an Illumina Hiseq 4000. For samples from donors 390c and 417c, sixteen 10x Genomics Chromium 5’ libraries were pooled and sequenced on 2 lanes of a S2 flowcell of Illumina Novaseq with 50bp paired end reads.

### 10x Genomics Chromium VDJ library preparation and sequencing

10x Genomics VDJ libraries were generated from the 5’ 10x Genomics Chromium cDNA libraries as detailed in the manufacturer’s protocol. BCR libraries for each sample were pooled and sequenced on a single lane of Illumina HiSeq 4000 with 150bp paired end reads.

### Plate-based scRNA-seq

Plate based scRNA-seq was performed with the NEBNext Single Cell/Low Input RNA Library Prep Kit for Illumina (New England Biolabs Inc, E6420L). Cells from donor 364b were snap frozen in 10% DMSO in 90% BSA. Cells were thawed rapidly in a 37°C water bath and diluted slowly with pre-warmed 2% FBS in D-PBS. Cells were pelleted for 5 minutes at 300g and washed with 500ul of DPBS and pelleted as before. Cells were resuspended in 200ul of CD25-PE (clone M-A251, Biolegend), CD127-FITC (clone EBIORDR5, eBioscience), CD4-BV421 (clone SK3, Biolegend) and TCRα/β-APC (clone 1p26, Biolegend) (all diluted 1:200) and Zombie Aqua Fixable Viability Dye (Biolegend) (diluted 1:1000) and incubated for 30 minutes in the dark at room temperature. Cells were washed twice with 500ul of 2% FBS in D-PBS before being filtered through a 100uM filter. Single, live, TCRβ+ cells were FACS sorted into a pre-prepared 384-well plate (Eppendorf, Cat. No. 0030128508) containing 2 *µ*l of 1X NEBNext Cell Lysis Buffer. FACS sorting was performed with a BD Influx sorter with the indexing setting enabled. Plates were sealed and spun at 100 x g for 1 minute then immediately frozen on dry ice and stored at −80°C.

cDNA generation was then performed in an automated manner on the Agilent Bravo NGS workstation (Agilent Technologies). Briefly, 1.6 *µ*l of Single Cell RT Primer Mix was added to each well and annealed on a PCR machine (MJ Research Peltier Thermal Cycler) at 70°C for 5 minutes. 4.4 *µ*l of Reverse Transcription (RT) mix was added to the mixture and further incubated at 42°C for 90 minutes followed by 70°C for 10 minutes to generate cDNA. 22 *µ*l of cDNA amplification mix containing NEBNext Single Cell cDNA PCR MasterMix and PCR primer was mixed with the cDNA, sealed and spun at 100 x g for 1 minute. cDNA amplification was then performed on a PCR machine (MJ Research Peltier Thermal Cycler) with 98°C 45 s, 20 cycles of [98°C 10 s, 62°C 15 s, 72°C 3 mins], 72°C 5 mins. The 384-well plate containing the amplified cDNA was purified with an AMPure XP workflow (Beckman Coulter, Cat No. A63880) and quantified with the Accuclear Ultra High Sensitivity dsDNA kit (Biotium, Cat. No. 31028). ∼10 ng of cDNA was stamped into a fresh 384-well plate (Eppendorf, Cat. No. 0030128508) for sequencing library preparation.

Sequencing libraries were then generated on the Agilent Bravo NGS workstation (Agilent Technologies). Purified cDNA was fragmented by the addition of 0.8 *µ*l of NEBNext Ultra II FS Enzyme Mix and 2.8 *µ*l of NEBNext Ultra II FS Reaction buffer to each well and incubated on a PCR machine (MJ Research Peltier Thermal Cycler) for 72°C at 15 minutes and 65°C for 30 minutes. A ligation mixture was then prepared containing NEBNext Ultra II Ligation Master Mix, NEBNext Ligation Enhancer and 100 *µ*M Illumina compatible adapters (Integrated DNA Technologies) and 13.4 *µ*l added to each well of the 384-well plate. The ligation reaction was incubated on the Agilent workstation at 20°C for 15 minutes and then purified and size selected with an AMPure XP workflow (Beckman Coulter, Cat No. A63880). 20 *µ*l of KAPA HiFi HS Ready Mix (Kapa Biosystems, Cat. No. 07958927001) was then added to a pre-prepared 384-well plate (Eppendorf, Cat. No. 0030128508) containing 100 *µ*M i5 and i7 indexing primer mix (50 *µ*M each) (Integrated DNA Technologies). The indexing primers pairs were unique to allow multiplexing of up to 384 single cells in one sequencing pool. The plate containing the PCR Master Mix and indexing primers was stamped onto the adapter ligated purified cDNA, sealed and spun at 100 x g for 1 minute. Amplification was performed on a PCR machine (MJ Research Peltier Thermal Cycler) with 95°C for 5 minutes, 8 cycles of [98°C 30 seconds, 65°C 30 seconds, 72°C 1 minute], 72°C 5 minutes. The PCR products were pooled in equal volume on the Microlab STAR automated liquid handler (Hamilton Robotics) and the pool purified and size selected with an AMPure XP workflow (Beckman Coulter, Cat No. A63880). The purified pool was quantified on an Agilent Bioanalyser (Agilent Technologies) and sequenced on one lane of an Illumina HiSeq 4000 instrument.

### Single-cell RNA sequencing data alignment

10x Genomics gene expression raw sequencing data was processed using CellRanger software version 3.0.2 and 10X human genome GRCh38-3.0.0 as the reference. 10x Genomics VDJ immunoglobulin heavy and light chain were processed using cellranger vdj (version 3.1.0) and the reference cellranger-vdj-GRCh38-alts-ensembl-3.1.0 with default settings. NEB sequencing data was processed using STAR 2.5.1b into HTSeq and mapped to 10X human genome GRCh38-1.2.0.

### Single-cell RNA sequencing quality control

Single cell read counts from all samples were pooled and filtered considering number of UMIs - keeping genes expressed in minimum of 3 cells, keeping cells where genes detected are in a range 700-6000. Non-immune cells were excluded from the final analysis based on the absence of PTPRC and presence of markers such as EPCAM (epithelial cells) and COL1A1 (fibroblasts).

### Cell type annotation

Cells were clustered using Scanpy (version 1.4) processing pipeline ^56^. In short, the counts were normalised to 10,000 reads per cell (sc.pp.normalise_per_cell) and log transformed (sc.pp.log1p) to be comparable amongst the cells. The number of UMIs and percentage of mitochondrial genes were regressed out (sc.pp.regress_out) and genes were scaled (sc.pp.scale) to unit variance. The normalised counts were used to detect highly variable genes (sc.pp.highly_variable_genes). Batch correction between the donors was performed using bbknn method ^57^ on 50 PCs and trim parameter set to 100. Clusters were then identified using Leiden graph-based clustering (resolution set to 1). Cell identity was assigned using known markers shown in Figure 1E and the top differentially expressed genes identified using Wilcoxon rank-sum test (sc.tl.rank_genes_groups function, Supplementary Table 4). CD4 T cells, myeloid cells, B cells were subclustered for identification of subsets within each cluster. Treg cells annotated above were further subclustered using the “FindClusters” and “RunUMAP” functions from Seurat (Version 3.0.1)^58^ (Figure 3).

The number of PCs used for Treg cell clustering were estimated by the elbow of a PCA screen plot, in combination to manual exploration of the top genes from each PC. Clustering of mLN Treg cells was performed using 1-14 PCs and resolution of 0.3 and colonic Treg cells with 1-11 PCs and resolution of 0.3.

### ‘Pseudospace’ analysis

‘Pseudospace’ analysis of Treg cells was performed using Monocle 2.2.0 ^59^. Data was log normalised and cell ordered based on DDRTree reduction on highly variable genes with donor effect regression. The heatmap in Figure 3c was generated using the “plot_pseudotime_heatmap” function in Monocle 2.2.0 ^59^. Genes contributing to ‘pseudospace’ were first filtered for exclusion of mitochondria and immunoglobulin genes, expression in at least 15 cells and qval < 0.001. Gene expression was smoothed into 100 bins along pseudospace using a natural spline with 3 degrees of freedom. Matching column annotation for each bin was determined as the most prevalent tissue region origin of the cells within that bin. Genes were grouped into 2 clusters and ordered by hierarchical clustering. Cells were ordered through pseudotime.

### Sampling of microbiome

Swabs were taken immediate of the mucosal surface of excised tissue using MWE Transwab Cary Blair (catalogue number: MW168). Swabs were maintained at 4°C. Working in a biosafety cabinet, swabs were washed in 500*µ*l of anaerobic PBS and mixed with anaerobic 50% glycerol before being snap frozen on dry ice and stored at −80°C until use.

### Microbiota profiling and sequencing

The microbiota vials were defrosted on ice. Approximately 100*µ*l of each was transferred into new Eppendorf tubes. DNA was extracted from microbiome samples using the MP Biomedical FastDNA SPIN Kit for soil (catalogue number 116560200). 16S rRNA gene amplicon libraries were made by PCR amplification of variable regions 1 and 2 of the 16S rRNA gene using the Q5 High-Fidelity Polymerase Kit supplied by New England Biolabs as described in ^60^. Primers 27F AATGATACGGCGACCACCGAGATCTACAC (first part, Illumina adaptor) TATGGTAATT (second part, forward primer pad) CC (third part, forward primer linker) AGMGTTYGATYMTGGCTCAG (fourth part, forward primer) and 338R CAAGCAGAAGACGGCATACGAGAT (first part, reverse complement of 3′ Illumina adaptor) ACGAGACTGATT (second part, golay barcode) AGTCAGTCAG (third part, reverse primer pad) AA (fourth part, reverse primer linker) GCTGCCTCCCGTAGGAGT (fifth part, reverse primer) were used. Four PCR amplification reactions per sample were carried out; products were pooled and combined in equimolar amounts for sequencing using the Illumina MiSeq platform, generating 150□bp reads. Analysis of partial 16S rRNA sequences was carried out using SILVA v132 and mothur MiSeq SOP^61^. The 16S rRNA gene alignments were used to determine a maximum likelihood phylogeny using Fasttree ^62^. Phylogenetic trees were visualised and edited using iTOL^63^.

### T cell clonal sharing analysis

T cell receptor sequences generated using the Smartseq2 scRNA-seq protocol were reconstructed using the TraCeR software as previously described ^64^.

### Bulk B cell receptor sequencing

Small portions of samples were taken from excised tissues and snap frozen in 1ml of RNAlater (Ambion). RNA extracted from tissue using the QIAshreddar and QIAGEN Mini Kit (50). RNA concentration was measured using a Bioanalyser. B Cell Receptor (BCR) heavy chain sequences of all B lineage subsets present in the tissue were amplified as previously described ^41^. Briefly, RNA was reverse transcribed using a barcoded reverse primer set capturing all antibody (sub)classes. Targeted heavy-chain amplification was performed with a multiplex set of IGHV gene primers to FR1 and a universal reverse primer using HiFI qPCR KAPA Biosystems. After adapter filtering and trimming, BCR sequences were assembled and aligned using MiXCR (version 65)^65^. It is worth noting that detected BCR sequences are biased towards those included in the reference database and while there is a continuous discovery of novel germline alleles, no database is currently a complete reflection of the human IGH locus diversity. Only in-frame and IGH sequences with at least 3 read counts were kept for the analysis. To calculate the CDR3 nucleotide shared repertoire, the tcR package was used ^66^.

### 10x Genomics single-cell VDJ data processing, quality control and annotation

Poor quality VDJ contigs that either did not map to immunoglobulin chains or were assigned incomplete by cellranger were discarded. For additional processing, all IgH sequence contigs per donor were combined together. We further filtered IgH contigs as to whether they had sufficient coverage of constant regions to ensure accurate isotype assignment between closely related subclasses using MaskPrimers.py (pRESTO version 0.5.10) ^67^. IgH sequences were then further annotated using IgBlast ^68^ and reassigned isotype classes using AssignGenes.py (pRESTO) prior to correction of ambiguous V gene assignments using TIgGER (version .03.1) ^69^.

Clonally-related IgH sequences were identified using DefineClones.py (ChangeO version 0.4.5) ^69^ with a nearest neighbour distance threshold of 0.2, as determined by visual inspection of the output of distToNearest (Shazam version 0.1.11) ^69^. CreateGermlines.py (ChangeO version 0.4.5) was then used to infer germline sequences for each clonal family and observedMutations (Shazam version 0.1.11) was used to calculate somatic hypermutation frequencies for each IgH contig. Estimated clonal abundances and IgH diversity analyses within each donor were performed using estimateAbundance, rarefyDiversity and testDiversity of Alakazam (version 0.2.11) ^69^ with a bootstrap number of 500. Finally, the number of quality filtered and annotated IgH, IgK or IgL were determined per unique cell barcode. If more than one contig per chain was identified, metadata for that cell was ascribed as “Multi”. The subsequent metadata table was then integrated with the single-cell RNA-seq gene expression objects for annotation of IgH contigs with B cell types and downstream analysis. Co-occurence of expanded clone members between tissues and/or cell types was reported as a binary event for each clone that contained a member within two different tissues and/or cell types.

### Quantifying IgA binding of bacteria

Microbiome frozen in 50% glycerol were defrosted on ice before being washed in PBS and pelleted for 3 minutes at 8,000 rpm. Bacteria were then stained on ice for 30 minutes with IgG-PE (Biolegend, clone HP6017), IgA1/2 biotin (BD, clone G20-359), followed by 20 minutes with streptavidin-APC (Biolegend). For isotype controls, mouse IgG1 κ Isotype-Biotin (BD) or mouse IgG2a, κ Isotype-PE (Biolegend) were used. The bacteria were washed before and after DNA was stained with Hoechst33342 (Sigma-Aldrich). The stained bacteria were sorted as Hoechst^+^ and IgG^+^IgA^+^ or IgG^-^IgA^+^ into PBS using the BD Influx and then stored at −80°C until DNA extraction.

### Genomic DNA Extraction for shotgun sequencing of IgA-bound bacteria

Bacteria in PBS was defrosted on ice before being pelleted at 3900 rpm for 5 minutes at 4°C. The supernatant was removed and the pellet resuspended in 2ml of 25% sucrose in TE Buffer (10mM Tris pH8 and 1mM EDTA pH8). 50*µ*l of 100mg/ml lysozyme in 0.25M Tris was added and incubated at 37°C for 1 hour. 100ul of Proteinase K (18mg/ml),15*µ*l of RNAase A (20mg/ml), 400*µ*l of 0.5M EDTA (pH 8), and 250*µ*l of 10% Sarkosyl were then added. This was left on ice for 2 hours and then incubated at 50°C. The DNA was then isolated by running through a gel, samples quantitated by qbit and pooled. Pooled DNA was sequenced on the Illumina Hiseq2500 platform. Inverse simpson (diversity) and chao (richness) of IgA-opsonised bacteria was determined by using R package “microbiome”.

## Data availability

Raw sequencing data files are available at ArrayExpress (accession numbers: E-MTAB-8007, E-MTAB-8474, E-MTAB-8476, E-MTAB-8484 & E-MTAB-8486). Sequencing data for the microbiome are available at MGnify (ERS numbers are listed in Supplementary Table 2). Processed single-cell RNA sequencing object is available for online visualisation and download at gutcellatlas.org.

## Figure Legends

**Supplementary Figure 1: Single-cell mapping of immune cells in the steady-state human colon.**

a) Shannon diversity score of bacterial OTUs from mucosa surfaces of the caecum, transverse colon and sigmoid colon (n=12 donors). b) Relative abundances of OTUs at phylum level as in a. c) Relative abundances of *Bacteroides, Coprobacillus, Shigella* and *Enterococcus spp*. in colon regions as in a, with lines connecting samples from each donor. d) Flow cytometry-sorting strategy for single, live, CD45^+^, CD3^-^, CD4^-^ immune cells (red) and single, live, CD45^+^, CD3^+^, CD4^+^ T cells (blue) from mLN (top) and colon lamina propria (bottom). e) UMAP projection of all cells after filtering and QC (as in Figure 1d & 1e), with the five donors represented as different colours. b & c) Bars show mean proportion in each region and statistical significances are calculated from 2-way ANOVA with Tukey’s multiple corrections of the total OTU dataset. * *P* <0.05; ** *P* <0.01; **** *P* <0.0001

**Supplementary Figure 2: Defining T helper cell transcriptional signatures in the colon.**

a) Heatmap of differentially expressed genes with filtering of 0.25 log fold change and minimum percent expression of 10%between Th1 and Th17 cells pooled from caecum, transverse colon and sigmoid colon (n=5 donors). Rows are ordered by hierarchical clustering and columns are grouped by cell type as denoted in the column annotation.

**Supplementary Figure 3: Treg subsets form a lymphoid-to-periphery activation trajectory.**

a) Log mean expression level and percentage of marker genes used to annotate Treg subsets in mLN and colon (n=5 donors). Gradient bar represents shift in function of marker genes. b) tSNE projection of clustering of Treg subsets and conventional CD4+ T cells of mLN and colon dataset.

**Supplementary Figure 4: Characterising B cell phenotypes and BCR expression in mLN and colon regions.**

a) BCR diversity rarefaction analysis of IgA+ plasma cell clones across multiple diversity orders (Q) using single-cell VDJ-derived IgH sequences (n=2 donors). This reveals comparable diversity between donors. b) Same as in a), but expanded IgA+ plasma cell clones from both donors were pooled and were instead separated by gut regions for BCR diversity rarefaction analysis (upper panel). Lower panels represent the results from significance tests of the diversity index between gut regions at either Q = 0 (left; corresponding to species richness measure) or Q = 1 (right; corresponding to the exponential of the Shannon-Weiner index). c) Co-occurrence of expanded B cell clones identified by single-cell VDJ analysis that are shared across different gut regions and between different cell types. Numbers reflect a binary detection event rather than the number of members per clone shared. d) Frequency of variable and joining BCR sequence expression in mLN and colon regions, detected by bulk BCR sequencing (n=3 donors). Rows are ordered by hierarchical clustering.

**Supplementary Table 1:** Metadata of deceased transplant donors involved in this study.

| ID | Sex | Age | Death | Approx. BMI | CMV/EBV | Antibiotics prescribed within 2 weeks of death |
| --- | --- | --- | --- | --- | --- | --- |
| 287c | M | 50-54 | DCD | 22 | CMV-/EBV+ | Ceftriaxone, Meropenem, Metronidazole |
| 290b | F | 60-64 | DBD | 28 | CMV+/EBV+ |  |
| 296c | F | 35-40 | DCD | 21 | CMV-/EBV+ | Tazocin |
| 298c | M | 55-60 | DCD | 24 | CMV-/EBV+ |  |
| 302c | M | 40-44 | DCD | 35 | CMV-/EBV- |  |
| 312c | M | 75-80 | DCD | 24 | CMV+/EBV+ |  |
| 362c | M | 60-65 | DCD | 27 | CMV-/EBV+ |  |
| 364b | M | 50-55 | DBD | 26 | CMV-/EBV+ |  |
| 367b | M | 65-70 | DBD | 25 | CMV+/EBV+ |  |
| 390c | F | 65-70 | DCD | 35 | CMV+/EBV+ |  |
| 403c | M | 50-55 | DCD | 33 | CMV+/EBV+ | Tazocin |
| 411c | M | 25-30 | DCD | 25 | CMV+/EBV+ | Meropenem |
| 414c | M | 50-55 | DCD | 30 | CMV-/EBV+ |  |
| 417c | M | 65-70 | DCD | 25 | CMV-/EBV+ |  |

## References

1. Halfvarson, J. et al. Dynamics of the human gut microbiome in inflammatory bowel disease. Nat Microbiol 2, 17004 (2017).

2. Donaldson, G. P., Melanie Lee, S. & Mazmanian, S. K. Gut biogeography of the bacterial microbiota. Nature Reviews Microbiology 14, 20–32 (2016).

3. Zoetendal, E. G. et al. The human small intestinal microbiota is driven by rapid uptake and conversion of simple carbohydrates. ISME J. 6, 1415–1426 (2012).

4. Cummings, J. H. & Macfarlane, G. T. The control and consequences of bacterial fermentation in the human colon. J. Appl. Bacteriol. 70, 443–459 (1991).

5. Wang, X., Heazlewood, S. P., Krause, D. O. & Florin, T. H. J. Molecular characterization of the microbial species that colonize human ileal and colonic mucosa by using 16S rDNA sequence analysis. J. Appl. Microbiol. 95, 508–520 (2003).

6. Mowat, A. M. & Agace, W. W. Regional specialization within the intestinal immune system. Nat. Rev. Immunol. 14, 667–685 (2014).

7. Denning, T. L. et al. Functional Specializations of Intestinal Dendritic Cell and Macrophage Subsets That Control Th17 and Regulatory T Cell Responses Are Dependent on the T Cell/APC Ratio, Source of Mouse Strain, and Regional Localization. The Journal of Immunology 187, 733–747 (2011).

8. Atarashi, K. et al. Th17 Cell Induction by Adhesion of Microbes to Intestinal Epithelial Cells. Cell 163, 367–380 (2015).

9. Ivanov, I. I. et al. Induction of intestinal Th17 cells by segmented filamentous bacteria. Cell 139, 485–498 (2009).

10. Atarashi, K. et al. Induction of colonic regulatory T cells by indigenous Clostridium species. Science 331, 337–341 (2011).

11. Mazmanian, S. K., Liu, C. H., Tzianabos, A. O. & Kasper, D. L. An immunomodulatory molecule of symbiotic bacteria directs maturation of the host immune system. Cell 122, 107–118 (2005).

12. Atarashi, K. et al. Ectopic colonization of oral bacteria in the intestine drives TH1 cell induction and inflammation. Science 358, 359–365 (2017).

13. Fanning, S. et al. Bifidobacterial surface-exopolysaccharide facilitates commensal-host interaction through immune modulation and pathogen protection. Proc. Natl. Acad. Sci. U. S. A. 109, 2108–2113 (2012).

14. Bozorov, T. A., Rasulov, B. A. & Zhang, D. Characterization of the gut microbiota of invasive Agrilus mali Matsumara (Coleoptera: Buprestidae) using high-throughput sequencing: uncovering plant cell-wall degrading bacteria. Sci. Rep. 9, 4923 (2019).

15. Consortium, T. H. M. P. & The Human Microbiome Project Consortium. Structure, function and diversity of the healthy human microbiome. Nature 486, 207–214 (2012).

16. Wu, G. D. et al. Linking long-term dietary patterns with gut microbial enterotypes. Science 334, 105–108 (2011).

17. Jones, R. B. et al. Inter-niche and inter-individual variation in gut microbial community assessment using stool, rectal swab, and mucosal samples. Sci. Rep. 8, 4139 (2018).

18. Smillie, C. S. et al. Rewiring of the cellular and inter-cellular landscape of the human colon during ulcerative colitis. (2018). doi: 10.1101/455451

19. Chakarov, S. et al. Two distinct interstitial macrophage populations coexist across tissues in specific subtissular niches. Science 363, (2019).

20. Kumar, B. V. et al. Human Tissue-Resident Memory T Cells Are Defined by Core Transcriptional and Functional Signatures in Lymphoid and Mucosal Sites. Cell Rep. 20, 2921–2934 (2017).

21. Cook, D. N. et al. CCR6 mediates dendritic cell localization, lymphocyte homeostasis, and immune responses in mucosal tissue. Immunity 12, 495–503 (2000).

22. Ward, S. Faculty of 1000 evaluation for Transcription factor KLF2 regulates the migration of naive T cells by restricting chemokine receptor expression patterns. F1000 - Post-publication peer review of the biomedical literature (2008). doi: 10.3410/f.1103864.560997

23. Toribio-Fernández, R. et al. Lamin A/C augments Th1 differentiation and response against vaccinia virus and Leishmania major. Cell Death Dis. 9, 9 (2018).

24. Mateyak, M. K. & Kinzy, T. G. eEF1A: thinking outside the ribosome. J. Biol. Chem. 285, 21209–21213 (2010).

25. Lönnberg, T. et al. Single-cell RNA-seq and computational analysis using temporal mixture modelling resolves Th1/Tfh fate bifurcation in malaria. Sci Immunol 2, (2017).

26. Miragaia, R. J. et al. Single-Cell Transcriptomics of Regulatory T Cells Reveals Trajectories of Tissue Adaptation. Immunity (2019). doi: 10.1016/j.immuni.2019.01.001

27. Miyao, T. et al. Plasticity of Foxp3 T Cells Reflects Promiscuous Foxp3 Expression in Conventional T Cells but Not Reprogramming of Regulatory T Cells. Immunity 36, 262–275 (2012).

28. Mucida, D. et al. Oral tolerance in the absence of naturally occurring Tregs. J. Clin. Invest. 115, 1923–1933 (2005).

29. Povoleri, G. A. M. et al. Human retinoic acid-regulated CD161 regulatory T cells support wound repair in intestinal mucosa. Nat. Immunol. 19, 1403–1414 (2018).

30. Schiering, C. et al. The alarmin IL-33 promotes regulatory T-cell function in the intestine. Nature 513, 564–568 (2014).

31. Krzysiek, R. et al. Antigen receptor engagement selectively induces macrophage inflammatory protein-1 alpha (MIP-1 alpha) and MIP-1 beta chemokine production in human B cells. J. Immunol. 162, 4455–4463 (1999).

32. Takahashi, K. et al. CCL3 and CCL4 are biomarkers for B cell receptor pathway activation and prognostic serum markers in diffuse large B cell lymphoma. British Journal of Haematology 171, 726–735 (2015).

33. Burger, J. A. et al. High-level expression of the T-cell chemokines CCL3 and CCL4 by chronic lymphocytic leukemia B cells in nurselike cell cocultures and after BCR stimulation. Blood 113, 3050–3058 (2009).

34. Mencarelli, A. et al. Highly specific blockade of CCR5 inhibits leukocyte trafficking and reduces mucosal inflammation in murine colitis. Sci. Rep. 6, 30802 (2016).

35. Zhang, Y. et al. Plasma cell output from germinal centers is regulated by signals from Tfh and stromal cells. J. Exp. Med. 215, 1227–1243 (2018).

36. Caraux, A. et al. Circulating human B and plasma cells. Age-associated changes in counts and detailed characterization of circulating normal CD138- and CD138 plasma cells. Haematologica 95, 1016–1020 (2010).

37. Hieshima, K. et al. CC chemokine ligands 25 and 28 play essential roles in intestinal extravasation of IgA antibody-secreting cells. J. Immunol. 173, 3668–3675 (2004).

38. Hu, K. et al. CCL19 and CCL28 Augment Mucosal and Systemic Immune Responses to HIV-1 gp140 by Mobilizing Responsive Immunocytes into Secondary Lymph Nodes and Mucosal Tissue. The Journal of Immunology 191, 1935–1947 (2013).

39. Mora, J. R. & von Andrian, U. H. Differentiation and homing of IgA-secreting cells. Mucosal Immunol. 1, 96 (2008).

40. Zhang, W. et al. Characterization of the B Cell Receptor Repertoire in the Intestinal Mucosa and of Tumor-Infiltrating Lymphocytes in Colorectal Adenoma and Carcinoma. The Journal of Immunology 198, 3719–3728 (2017).

41. Petrova, V. N. et al. Combined Influence of B-Cell Receptor Rearrangement and Somatic Hypermutation on B-Cell Class-Switch Fate in Health and in Chronic Lymphocytic Leukemia. Front. Immunol. 9, 1784 (2018).

42. Macpherson, A. J. A Primitive T Cell-Independent Mechanism of Intestinal Mucosal IgA Responses to Commensal Bacteria. Science 288, 2222–2226 (2000).

43. Harbour, S. N., Maynard, C. L., Zindl, C. L., Schoeb, T. R. & Weaver, C. T. Th17 cells give rise to Th1 cells that are required for the pathogenesis of colitis. Proc. Natl. Acad. Sci. U. S. A. 112, 7061–7066 (2015).

44. Ferreira, R. C. et al. Cells with Treg-specific FOXP3 demethylation but low CD25 are prevalent in autoimmunity. J. Autoimmun. 84, 75–86 (2017).

45. Hoffmann, P. et al. Loss of FOXP3 expression in natural human CD4+CD25+ regulatory T cells upon repetitive in vitro stimulation. Eur. J. Immunol. 39, 1088–1097 (2009).

46. Kyewski, B. & Suri-Payer, E. CD4+CD25+ Regulatory T Cells: Origin, Function and Therapeutic Potential. (Springer Science & Business Media, 2006).

47. Brandtzaeg, P. Function of mucosa-associated lymphoid tissue in antibody formation. Immunol. Invest. 39, 303–355 (2010).

48. Meng, W. et al. An atlas of B-cell clonal distribution in the human body. Nat. Biotechnol. 35, 879–884 (2017).

49. Dunn-Walters, D. K., Boursier, L. & Spencer, J. Hypermutation, diversity and dissemination of human intestinal lamina propria plasma cells. Eur. J. Immunol. 27, 2959–2964 (1997).

50. Tsuji, M. et al. Requirement for Lymphoid Tissue-Inducer Cells in Isolated Follicle Formation and T Cell-Independent Immunoglobulin A Generation in the Gut. Immunity 29, 261–271 (2008).

51. van der Waaij, L. A., Limburg, P. C., Mesander, G. & van der Waaij, D. In vivo IgA coating of anaerobic bacteria in human faeces. Gut 38, 348–354 (1996).

52. Imam, T., Park, S., Kaplan, M. H. & Olson, M. R. Effector T Helper Cell Subsets in Inflammatory Bowel Diseases. Front. Immunol. 9, 1212 (2018).

53. Castro-Dopico, T. et al. Anti-commensal IgG Drives Intestinal Inflammation and Type 17 Immunity in Ulcerative Colitis. Immunity (2019). doi: 10.1016/j.immuni.2019.02.006

54. Cunningham-Rundles, C. Physiology of IgA and IgA deficiency. J. Clin. Immunol. 21, 303–309 (2001).

55. Moon, C. et al. Vertically transmitted faecal IgA levels determine extra-chromosomal phenotypic variation. Nature 521, 90–93 (2015).

56. Wolf, F. A., Angerer, P. & Theis, F. J. SCANPY: large-scale single-cell gene expression data analysis. Genome Biol. 19, 15 (2018).

57. Park, J.-E., Polański, K., Meyer, K. & Teichmann, S. A. Fast Batch Alignment of Single Cell Transcriptomes Unifies Multiple Mouse Cell Atlases into an Integrated Landscape. doi: 10.1101/397042

58. Stuart, T. et al. Comprehensive Integration of Single-Cell Data. Cell 177, 1888–1902.e21 (2019).

59. Trapnell, C. et al. The dynamics and regulators of cell fate decisions are revealed by pseudotemporal ordering of single cells. Nature Biotechnology 32, 381–386 (2014).

60. Browne, H. P. et al. Culturing of ‘unculturable’ human microbiota reveals novel taxa and extensive sporulation. Nature 533, 543–546 (2016).

61. Kozich, J. J., Westcott, S. L., Baxter, N. T., Highlander, S. K. & Schloss, P. D. Development of a dual-index sequencing strategy and curation pipeline for analyzing amplicon sequence data on the MiSeq Illumina sequencing platform. Appl. Environ. Microbiol. 79, 5112–5120 (2013).

62. Price, M. N., Dehal, P. S. & Arkin, A. P. FastTree 2 – Approximately Maximum-Likelihood Trees for Large Alignments. PLoS One 5, e9490 (2010).

63. Letunic, I. & Bork, P. Interactive tree of life (iTOL) v3: an online tool for the display and annotation of phylogenetic and other trees. Nucleic Acids Res. 44, W242–5 (2016).

64. Stubbington, M. J. T. et al. T cell fate and clonality inference from single-cell transcriptomes. Nat. Methods 13, 329–332 (2016).

65. Bolotin, D. A. et al. MiXCR: software for comprehensive adaptive immunity profiling. Nat. Methods 12, 380–381 (2015).

66. Nazarov, V. I. et al. tcR: an R package for T cell receptor repertoire advanced data analysis. BMC Bioinformatics 16, (2015).

67. Heiden, J. A. V. et al. pRESTO: a toolkit for processing high-throughput sequencing raw reads of lymphocyte receptor repertoires. Bioinformatics 30, 1930–1932 (2014).

68. Ye, J., Ma, N., Madden, T. L. & Ostell, J. M. IgBLAST: an immunoglobulin variable domain sequence analysis tool. Nucleic Acids Research 41, W34–W40 (2013).

69. Gupta, N. T. et al. Change-O: a toolkit for analyzing large-scale B cell immunoglobulin repertoire sequencing data. Bioinformatics 31, 3356–3358 (2015).

