## Supplementary figures and images for "Distinct microbial and immune niches of the human colon"

### Supplementary Figures

#### Supp Figure 1

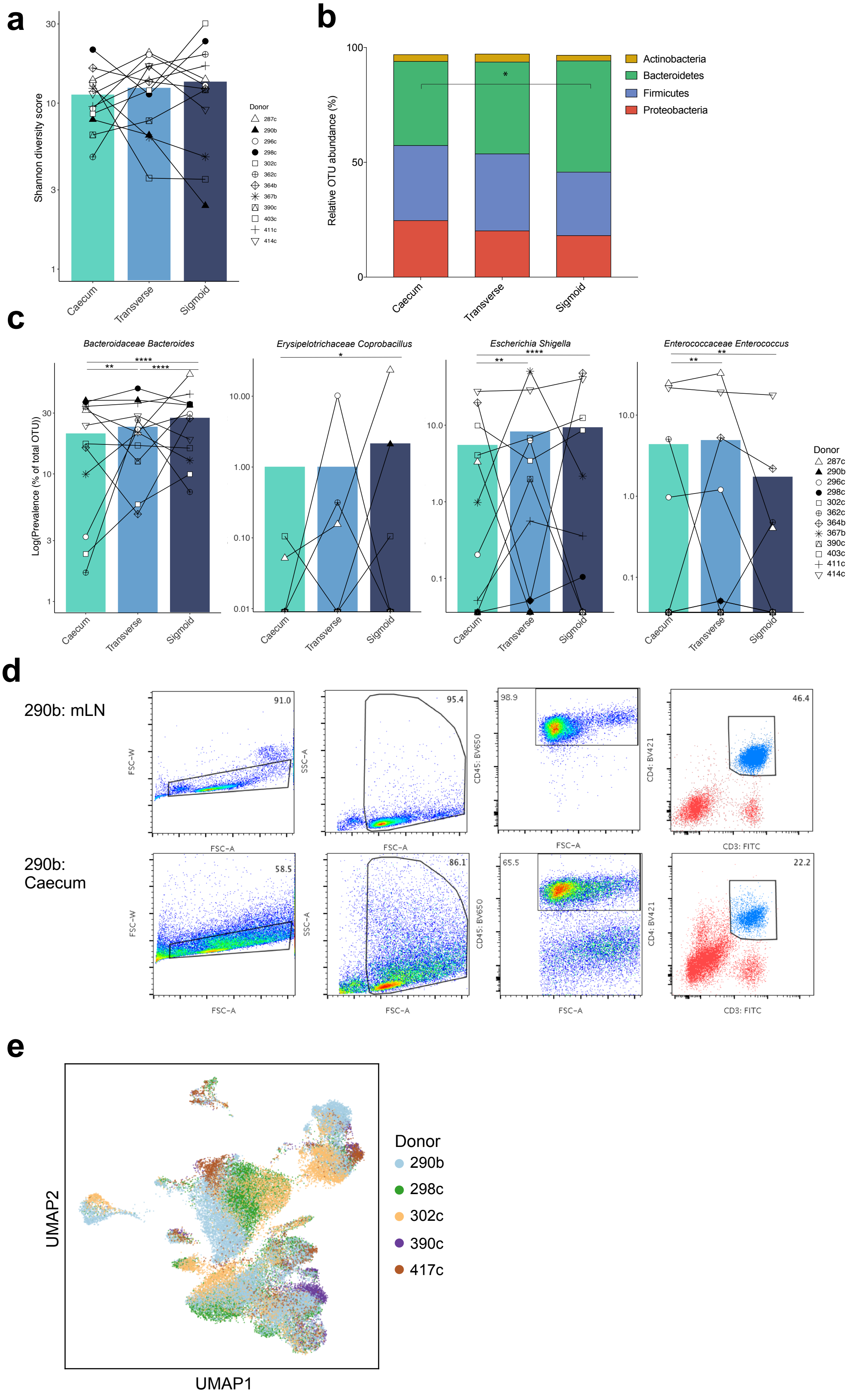

##### Supp Figure 2

**a**

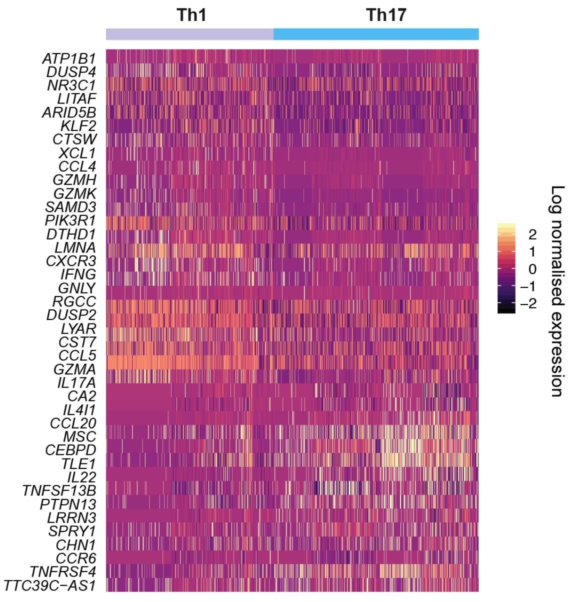

#### Supp Figure 3

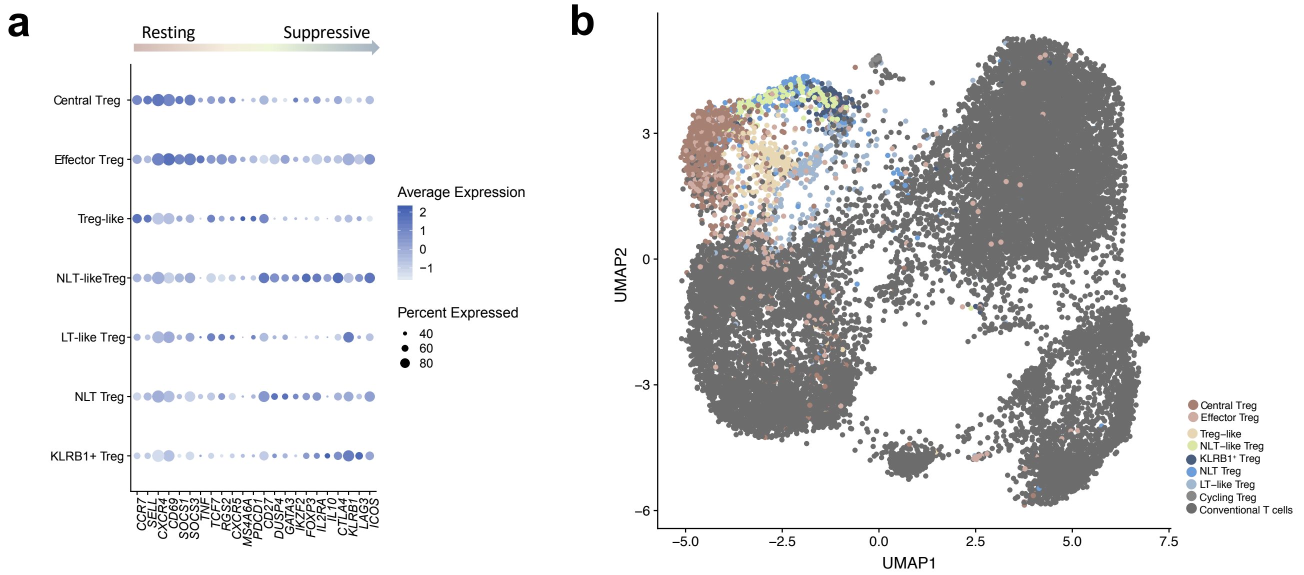

#### Supp Figure 4

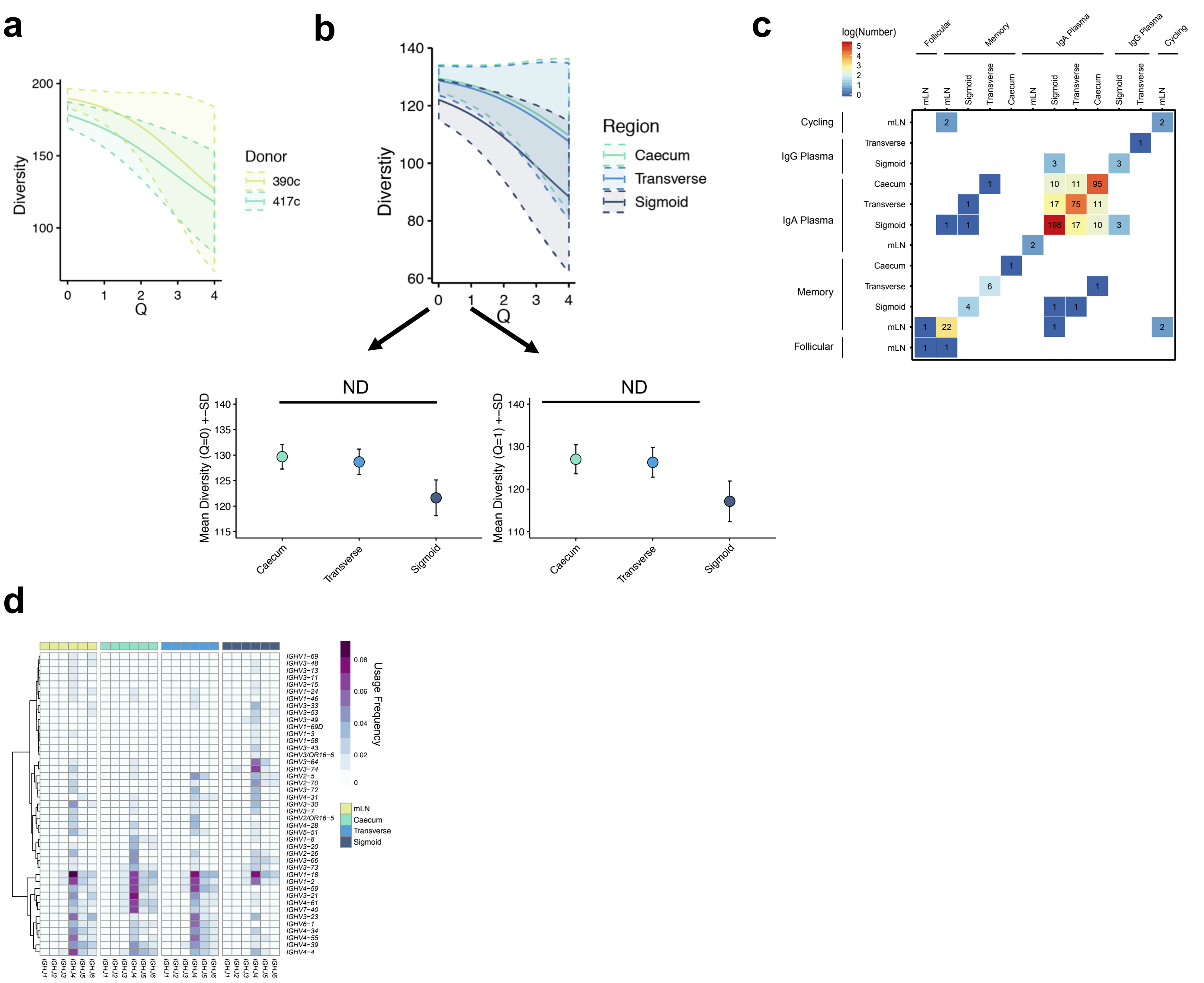
